# Within-population sperm competition intensity does not predict asymmetry in conpopulation sperm precedence

**DOI:** 10.1101/2020.04.23.055806

**Authors:** Martin D. Garlovsky, Leeban H. Yusuf, Michael G. Ritchie, Rhonda R. Snook

## Abstract

Postcopulatory sexual selection can generate coevolutionary arms races between the sexes resulting in the rapid coevolution of reproductive phenotypes. As traits affecting fertilisation success diverge between populations postmating prezygotic barriers to gene flow may evolve. Conspecific sperm precedence is a form of such isolation thought to evolve early during speciation yet has mostly been studied between species. Here we show conpopulation sperm precedence between *Drosophila montana* populations. Using genomic data to estimate divergence times and patterns of gene flow between populations, we show gene flow has played a considerable role during divergence. We find conpopulation sperm precedence is asymmetric and is concordant with asymmetry in non-competitive postmating prezygotic reproductive isolation. These results suggest these phenomena have a shared mechanism, but we show that this asymmetry is unrelated to the strength of postcopulatory sexual selection acting within populations. We tested whether overlapping foreign and coevolved ejaculates within the female reproductive tract altered fertilisation success but found no effect. Our results show that neither time since divergence nor sperm competitiveness predicts the strength of postmating prezygotic reproductive isolation. We suggest that divergence of postcopulatory phenotypes resulting in postmating prezygotic isolation is potentially driven by cryptic female choice, or mutation order divergence.

## INTRODUCTION

Widespread polyandry in animals presents the opportunity for postcopulatory sexual selection (sperm competition and cryptic female choice) and sexual antagonism to accelerate the coevolution of male ejaculate x female reproductive interactions [1–5]. Accordingly, reproductive traits such as gamete cell surface proteins, male seminal fluid proteins, female reproductive tract and sperm morphologies, show elevated rates of molecular and morphological evolution [6–9]. Barriers to gene flow between populations caused by reproductive interactions during or after mating but before fertilisation (i.e., postmating prezygotic, PMPZ) are expected to emerge early during speciation due to the rapid codiversification of reproductive traits within populations [10].

Identifying the barriers to gene flow that emerge earliest during reproductive isolation is key to understanding the origin of species [11–13]. Conspecific sperm precedence (CSP) is a widely observed form of competitive PMPZ isolation found where paternity is biased towards conspecifics when a female mates with both a con- and hetero-specific male [14,15]. CSP can result from postcopulatory selection either via conspecific sperm being more successful in sperm competition and/or favoured by cryptic female choice [5,16]. If postcopulatory sexual selection can facilitate the evolution of PMPZ isolation, then asymmetries in the strength of PMPZ isolation acting between taxa are predicted to reflect differences in the strength of sexual selection acting within populations. For instance, *in vitro* experiments in mouse (*Mus spp.*) have shown that CSP between taxa is correlated with the intensity of sperm competition acting within taxa [17,18]. No study has directly measured male paternity share in a competitive mating scenario *in vivo* and its relation to differences between populations in the strength of sperm competition experienced by males within populations. A pattern of CSP can also arise from direct incompatibility between the male ejaculate and female reproductive tract preventing heterospecific fertilisation [19]. Where females remate, overlapping coevolved and foreign ejaculates within the female reproductive tract might alter PMPZ outcomes [20]. For instance, a foreign male ejaculate could negatively affect the fertilisation function of a coevolved male ejaculate, or a coevolved ejaculate could provide the proper postmating female response improving fertility of a foreign male ejaculate [21].

CSP is thought to evolve early during speciation [22,23]. In sympatry CSP can evolve as a reinforcing mechanism [24–27]. The factors shaping the evolution of CSP during, as opposed to after, divergence remain poorly understood. Between divergent allopatric populations, conpopulation sperm precedence (CpSP) has been shown in some species [28,29], while other studies show little or no evidence of CpSP [17,30–32]. The demographic histories of taxa in studies of conpopulation sperm precedence are often poorly resolved, hampering inference about the evolution of reproductive isolation. For instance, it is not currently known whether conpopulation sperm precedence can evolve during divergence with gene flow.

We have previously described prezygotic reproductive isolation between three populations of *Drosophila montana*, from Crested Butte, Colorado, USA (referred to as Colorado), Oulanka, Finland, and Vancouver, Canada (Fig. S1) [33–37]. All three populations show premating and PMPZ isolation (Fig. S1). If reproductive isolation is solely due to isolation by distance, then the more distant Finnish population should be most divergent from the two North American populations and exhibit stronger reproductive isolation. However, total reproductive isolation is strongest between the geographically closer Colorado and Vancouver populations [35]. Vancouver females discriminate against Colorado males and both populations show non-competitive PMPZ isolation, where females successfully store sperm from foreign males after mating but lay many unfertilised eggs. PMPZ isolation is strongest when Colorado females mate with Vancouver males [35,36].

Using the *D. montana* system, we set out to ask (1) what is the relative timescale of divergence and what role has gene flow played during divergence, (2) do populations show conpopulation sperm precedence (CpSP) and if so, is it concordant with non-competitive PMPZ isolation, (3) is the strength of PMPZ isolation predicted by the strength of postcopulatory sexual selection acting within populations, and (4) do overlapping foreign and coevolved ejaculates interact to alter PMPZ outcomes? Previous estimates of population divergence were based on a few mtDNA and microsatellite markers [37]. To gain more accurate estimates of divergence time and the extent and time scale of ongoing or historical gene flow, we performed explicit demographic modelling of the focal populations using whole genome Pool sequencing data [38] and calculated within population genetic diversity. If CpSP is present between populations, and results from direct incompatibilities between the male ejaculate and female reproductive tract, then CpSP should be stronger in Colorado females than Vancouver [35,36]. We measured CpSP as male offensive paternity share (P2) when competing in a foreign or coevolved female reproductive tract against a foreign or coevolved male. We tested the prediction that PMPZ isolation asymmetries would reflect differences in the intensity of postcopulatory sexual selection acting within populations [18] by measuring two common proxies for the intensity of sperm competition faced by males; male reproductive mass investment and sequential mating capacity [39–41]. Both traits have been shown to be under sexual selection in other *Drosophila* species [42,43]. Finally, we tested the prediction that overlapping foreign and coevolved ejaculates within the female reproductive tract would alter PMPZ outcomes by calculating whether hatching success after a double mating differed from the expected additive effect of two single matings, given the estimated paternity share of the two males.

## METHODS

### Fly stocks

Wild mated female *Drosophila montana* were collected from riparian habitats in Crested Butte, Colorado, USA (38º49’N, 107º04’W), referred to as Colorado, Oulanka, Finland (66º22’N, 29º20’E), and Vancouver, Canada (48º55’N, 123º48’W). Population cages were established by combining 20 F3 males and females from each of 20 isofemale lines (800 flies total per population) in 2008 (Oulanka and Vancouver) and 2013 (Colorado population used for measuring reproductive isolation and reproductive investment) [35,36]. The Colorado population cage used for sequencing was established in 2009 from 13 isofemale lines (520 flies total) [35]. All stocks were maintained in large outbred populations on Lakovaara malt media [44] in overlapping generations in constant light at 19ºC to prevent females entering reproductive diapause. Flies were collected within three days of eclosion and kept in single sex vials of 10-20 individuals until reproductive maturity at 21-28 days old.

### Demographic modelling

#### Sequencing information

We obtained trimmed pool-seq data from Parker et al. [45]. Sequencing was performed in 2013, 5 years after collection for Oulanka and Vancouver, and 6 years for Colorado (around 50 generations). Genomic DNA was extracted from pooled samples of 50 females per population and sequenced using an Illumina HiSeq 2000 to produce paired-end reads (90 + 90 bp, insert size = 500 bp) at the Beijing Genomics Institute [45].

#### Sequence alignment and allele frequency estimation

We indexed the *D. montana* reference genome and mapped pool-seq reads using BWA-MEM v.0.75 [46]. We sorted reads, removed duplicates, and converted mapped reads to mpileup format using SAMtools v.1.6 [47] and converted this to a synchronised file using Popoolation2 [48]. All single nucleotide polymorphisms (SNPs) were filtered using a minimum coverage of 30 via the mpileip2sync.pl script. To account for fluctuations in coverage and allele frequency across loci, we subsampled SNPs to conform to a uniform coverage of 30 using the subsample-synchronized.pl script. To convert read counts into a format suitable for ∂a∂I (Diffusion Approximation for Demographic Inference) [38], we used the ‘poolfstat’ R package [49] and custom scripts.

We generated three-dimensional joint-site allele-frequency spectra (3D-AFS) and compared the fit of the spectra against competing demographic models using ∂a∂I using an existing pipeline [50]. To satisfy the assumption of independence between SNPs, we down-sampled from 860,228 to 8,000 SNPs by selecting 1 in every 100 SNPs.

#### Description of demographic models explored

We tested seven demographic models for the history of these populations (Table 1; Fig. S2). We fitted all seven models to the 3D-AFS and performed 3 rounds of model optimizations using the Nelder-Mead method (see supplementary material) [51]. We estimated log-likelihoods using a multinomial approach and performed model evaluation by calculating Akaike Information Criterion (AIC) for each model run [38].

**Table 1.** Results from demographic modelling using ∂a∂i comparing the 3D-allele frequency spectra on a dataset of 8,000 SNPs (see also Fig. 1).

| Model | log-lik | AIC | $\theta$ | nuOul | nuAm | nuVan | nuCol | mA | m1 | m2 | m3 | T1(a) | T1b | T2 | T3 |
| --- | --- | --- | --- | --- | --- | --- | --- | --- | --- | --- | --- | --- | --- | --- | --- |
| (1) No-migration | -24429.7 | 48871.4 | 194.07 | 0.170 | 4.439 | 5.392 | 4.522 |  |  |  |  | 3.525 |  | 3.106 |  |
| (2) Full symmetrical migration | -8923.31 | 17866.62 | 1397.29 | 0.243 | 0.473 | 0.237 | 0.551 | 0.317 | 0.785 | 4.486 | 0.603 | 0.875 |  | 11.810 |  |
| (3) Adjacent migration | -8668.5 | 17355 | 201.46 | 2.366 | 0.686 | 1.889 | 2.435 | 4.542 | 0.126 | 0.771 |  | 0.490 |  | 25.528 |  |
| (4) Period of ancient migration after initial divergence between North American populations and Oulanka | -9444.58 | 18903.16 | 1326.13 | 0.342 | 0.550 | 0.601 | 0.657 | 0.629 |  |  |  | 1.753 |  | 0.086 |  |
| (5) Ancient migration persisting after divergence between Colorado and Vancouver populations | -9931.74 | 19879.48 | 538.23 | 1.249 | 1.017 | 2.080 | 2.058 | 1.535 |  |  |  | 9.290 | 0.163 | 0.224 |  |
| (6) Short period of secondary contact beginning after divergence and a period of isolation between Colorado and Vancouver | -9285.95 | 18589.9 | 420.16 | 1.269 | 0.372 | 1.372 | 0.948 |  | 0.235 | 2.109 |  | 3.487 |  | 0.487 | 3.186 |
| (7) Secondary contact beginning during divergence between Colorado and Vancouver | -8747.33 | 17510.66 | 363.67 | 0.742 | 0.182 | 0.913 | 1.396 |  | 0.336 | 1.307 |  | 26.891 |  | 9.654 |  |

log-lik, log likelihood; AIC, Akaike Information Criterion; $\theta$ , nucleotide diversity; nuOul, nuVan, nuCol, effective population size in Oulanka, Vancouver, and Colorado, respectively; nuAm effective population size of precursor North American population; mA, ancient migration between Oulanka and precursor North American population; m1, ancient migration between Vancouver and Oulanka; m2, ancient migration between Vancouver and Colorado; m3, ancient migration between Oulanka and Colorado; Initial and secondary divergence are represented by T1(a) and T2, where T1(b) and T3 correspond to the start or end of periods of isolation. Population splits are denoted by T1 (between Oulanka and North America) and T2 (between Vancouver and Colorado). T1(a) and T1(b) denote the start or end of periods of migration following the split between Oulanka and North American populations. T3 denotes start or end of periods of migration following the split between Vancouver and Colorado.

#### Estimating within population genetic diversity

We mapped reads for each population to an updated *D. montana* genome (Poikela et al. in prep.) as above using BWA-MEM and conversion to pileup using SAMtools. To estimate θ_watterson_, θ_*π*_ and Tajima’s D, we used NPStat v.1 [52] and calculated summary statistics in 1kb windows, with minimum coverage (10), maximum coverage (100), minimum base quality (30) and minimum allele count (2) filters. We averaged windows for each scaffold to obtain scaffold-wide estimates of summary statistics and evaluated differences between populations using Kruskal-Wallis rank sum tests followed by pairwise Wilcox rank sum tests corrected for multiple testing using the Benjamini-Hochberg method.

### Measuring postmating prezygotic isolation

We measured postmating prezygotic isolation between Colorado and Vancouver using hatching success rates (a proxy for fertilisation success) [35,36] in single and double matings, and assessed male offensive paternity share (P2) using the irradiated male technique (see supplementary material) [53]. We performed experiments in 3 blocks for each focal female population separately, in which virgin females (n = 10 per cross-type per treatment per block) were presented with a virgin male from Colorado or Vancouver and allowed a 4-hour mating opportunity. After mating, males were discarded and females were transferred to a chamber in an oviposition manifold, which allows easy counting of numbers of eggs oviposited and assessment of hatching [35,36]. Manifolds were returned to the incubator for oviposition. Two days later, females were removed from their chambers, and placed in a vial with a second virgin male from Colorado or Vancouver and allowed a 4-hour remating opportunity. Remating females were returned to their chambers, placed back into the incubator and allowed to oviposit for a further two days on a new oviposition plate. We ensured no females mated more than once during either the first or second observation period. To measure hatching success the numbers of eggs laid on each oviposition plate were counted immediately after removing the female (for remating or discarding). The oviposition plate was then returned to the incubator for a further two days after which the numbers of unhatched eggs was counted again.

We assessed differences between cross-types in hatching success rates after the first and second mating using generalized linear models (GLMs) with binomial errors and a logit link, where the numbers of hatched (“successes”) and unhatched (“failures”) eggs was the response variable and the only predictor was the cross-type. We assessed differences in P2 using GLMs with binomial errors and a logit link where the binary response was the numbers of offspring sired by the second male (“successes”) and numbers of offspring sired by the first male (“failures”) (see supplementary material). Models included cross-type, irradiation order and their interaction as predictors. Females that did not remate, died prior to remating, or laid less than 10 eggs were excluded. Due to the low fertility in crosses between Colorado females and Vancouver males, we could not estimate P2 in the CVV cross, which was subsequently excluded from analyses (see supplementary material). We analysed responses in Colorado and Vancouver females separately as populations were never tested together. All statistical analyses were performed in R v.3.5.1 [54]. We used quasi-errors for GLMs where preliminary model inspection indicated overdispersion. Where appropriate we performed post-hoc Tukey’s honest significant difference (HSD) tests using the *glht* function from the ‘multcomp’ package [55].

#### Interaction between coevolved and foreign male ejaculates in the female reproductive tract

We tested whether observed hatching success rates after a double mating differed from the expected additive effect of two single matings, H_total_, using the equation:

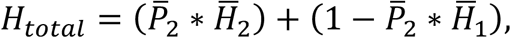

where 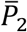 is the mean proportion of offspring sired by the second male in a given cross-type, and 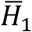 and 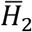 are the mean hatching success after a single mating for a female mated with a male from the first, and second, population denoted in a cross-type, respectively.

### Proxies measuring the intensity of sperm competition within populations

#### Male relative reproductive investment

Approximately 100 flies of mixed sex were collected from population cages and allowed to mate and oviposit for two days in plastic bottles covered with a molasses-agar oviposition plate and a drop of dried yeast paste added. Controlled density vials (CDVs) containing food were seeded with 50 first instar larvae from oviposition plates. Adults were collected from CDVs on the day of eclosion and kept in single sex vials until sexually mature and then frozen at −20ºC. Population identity was replaced with a unique identifier prior to measurement. Males (n = 60 per population) were thawed and the entire reproductive tract (testes, accessory glands, ejaculatory duct and bulb) dissected on a pre-weighed piece of foil in a drop of dH20 which was then transferred to a second piece of preweighed foil. Carcasses and reproductive tracts were dried overnight at 60ºC before weighing (METTLER TOLEDO^®^ UMX2 ultra-microbalance). We analysed male relative reproductive investment using analysis of covariance (ANCOVA), with log transformed dry reproductive tract mass as the response variable, and population ID (Colorado or Vancouver), log transformed dry soma mass (body mass – reproductive tract mass), and their interaction as predictors [39]. After removing four outliers (2 Colorado, 2 Vancouver; see supplementary material) we calculated ANOVA tables with type 2 sum of squares [56].

#### Male mating capacity

Virgin males (n = 20 per population) were mouth aspirated without anaesthesia into a vial containing two virgin females from their own population. Once copulation began unmated females were removed from the vial. The male was transferred to a new vial housing another two virgin females after mating ad libitum and we recorded the total number of matings that males performed within a 4-hour period. Mated females were returned to the incubator in their oviposition vial for 4 days, before being transferred to a second food vial for a further 4 days. The total number of adult flies emerging was counted for each female (combined across both oviposition vials). We tested for differences between populations in male sequential mating capacity and the total numbers of progeny sired by males across all their mates with Poisson GLMs. To test for differences between populations in male per mating investment we fitted a zero-inflated Poisson GLM using the ‘glmmTMB’ package [57]. Offspring numbers were used as the response variable, with population, mating number, and their interaction as predictors and male ID as a random effect. Two males were lost during the experiment (one Colorado, one Vancouver) and subsequently excluded from analyses.

#### Female dry mass

We measured dry mass of females as body size is known to correlate with fecundity in *Drosophila* [58]. Females emerging from CDVs (n = 60 per population) were frozen at −20ºC and later thawed and dried overnight at 60ºC before being weighed individually on a weighing boat.

## RESULTS

### Demographic modelling

Demographic analyses showed that the best fitting model included symmetric migration between Oulanka and Vancouver and between Colorado and Vancouver after splitting from the ancestral population suggesting a demographic history with substantial gene flow between populations (Fig. 1; Table 1; Fig. S3). The split between Colorado and Vancouver occurred shortly after the North American populations separated from Oulanka (Fig. 1). Additionally, Colorado and Oulanka showed larger effective population sizes than Vancouver (Table 1). However, ancestral population size for the North American population before the split was considerably lower, perhaps owing to population contraction following invasion of North America. For the best-fitting model, parameter estimates show considerably higher migration between the ancestral population and Oulanka (before Colorado and Vancouver split), than between Colorado and Vancouver or Vancouver and Oulanka (Fig. 1; Table 1).

**Figure 1.**
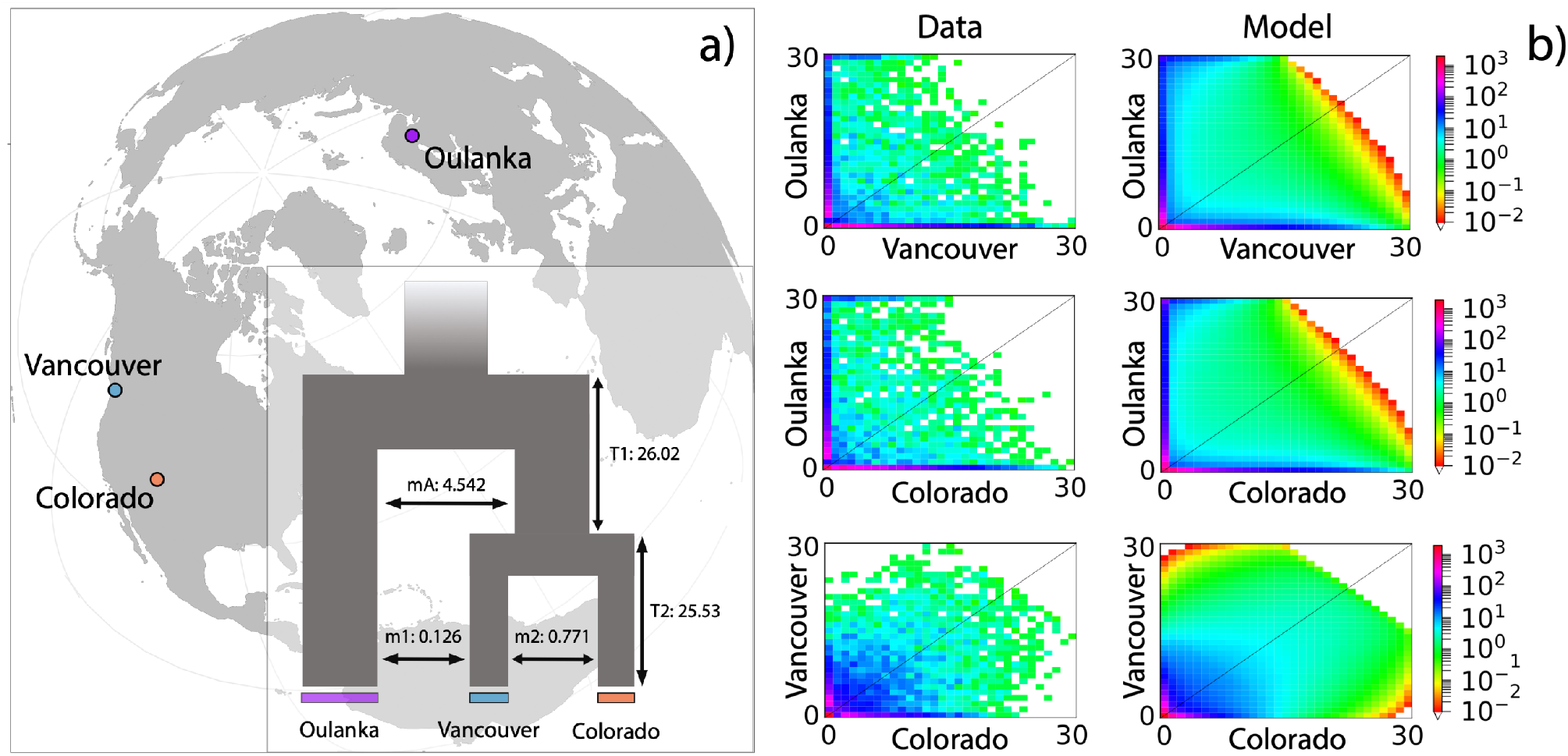
a) locations of *D. montana* populations. Inset: graphical representation of best fit model; adjacent migration between populations (Table 1). mA, ancient migration between Oulanka and precursor North American population; m1, ancient migration between Vancouver and Oulanka; m2, ancient migration between Vancouver and Colorado; T1, divergence between Oulanka and precursor North American population; T2, divergence between Colorado and Vancouver. b) 3D-allele frequency spectra of the empirical data (left) and the best model fit (right) in each pairwise combination between populations. Axes show counts of alleles in each population. Legends indicate number of sites in each cell. See Fig. S3 for plots including residuals.

For detailed description of models tested see supplementary material.

#### Within population genetic diversity

Tajima’s D (Kruskal-Wallace test, χ^2^ = 57.679, df = 2, p < 0.001) and pi (χ^2^= 7.38, df = 2, p = 0.025) both showed significant differences between populations but not Watterson’s theta (χ^2^ = 4.22, df = 2, p = 0.121) (Fig. 2). All populations showed negative genome-wide Tajima’s D, with Colorado and Oulanka exhibiting higher excesses of rare alleles than Vancouver, perhaps owing to recent changes in demography history (e.g. expansion following bottleneck), or selective sweeps (Fig. 2). This is supported by higher effective population size estimates in Colorado and Oulanka compared to Vancouver for the best-fitting demographic model (Table 1). Pi was significantly greater in Colorado than Oulanka (Wilcoxon rank sum test, p = 0.023), suggesting higher genetic diversity in Colorado than Oulanka (Fig. 2).

**Figure 2.**
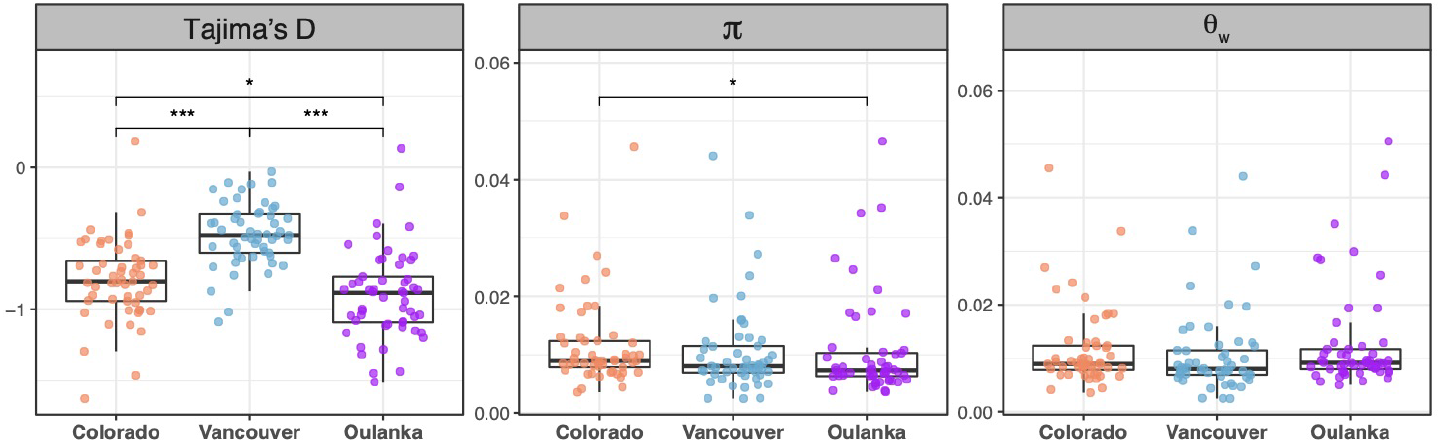
Genome-wide estimates of Tajima’s D, pi (*π*), and Watterson’s theta (θ_w_) calculated in 1kb windows. Points show the mean for each scaffold. Pairwise differences between populations were evaluated using Wilcox rank sum tests corrected for multiple testing using the Benjamini-Hochberg method: * p < 0.05, ** p < 0.01, *** p < 0.001.

### Postmating prezygotic reproductive isolation

#### Postmating prezygotic isolation of virgin females (non-competitive gametic isolation)

After a single mating, hatching success rates were similar to those reported previously [35,36]. Colorado females had reduced hatching success when mating with Vancouver males relative to mating with Colorado males (quasibinomial GLM: F_1,165_ = 341.97, p < 0.001, β = −3.89 ± 0.28). Likewise, Vancouver females had reduced hatching success when mating with Colorado males relative to mating with Vancouver males (quasibinomial GLM: F_1,136_ = 48.06, p < 0.001, β = −1.53 ± 0.23) (Fig. S4).

#### Postmating prezygotic isolation of non-virgin females

##### Hatching success rates

Cross-type had a significant effect on the proportion of eggs that hatched for Colorado females (quasibinomial GLM: F_3,111_ = 92.03, p < 0.001) and Vancouver females (quasibinomial GLM: F3,i37 = 34.86, p < 0.001) after the second mating (Fig. 3; Fig. S5). In both populations, females mating with a between-population male followed by a within-population male had hatching success not significantly different from those females mating with two within-population males (Tukey’s HSD: all p > 0.133; mean proportion of eggs that hatched: CCC = 0.84 ± 0.02, n = 26; CVC = 0.76 ± 0.03, n = 41; VVV = 0.88 ± 0.03, n = 29; VCV = 0.89 ± 0.02, n = 36). Females mating with two between-population males had lower hatching success than other groups (Tukey’s HSD: all p < 0.001; CVV = 0.12 ± 0.03, n = 30; VCC = 0.51 ± 0.03, n = 36). Females mating with a within-population male followed by abetween-population male had lower hatching success than females mating with two within-population males, but higher hatching success than females mating with two between-population males (Tukey’s HSD: all p < 0.016; CCV = 0.58 ± 0.06, n = 18; VVC = 0.72 ± 0.03, n = 40).

**Figure 3.**
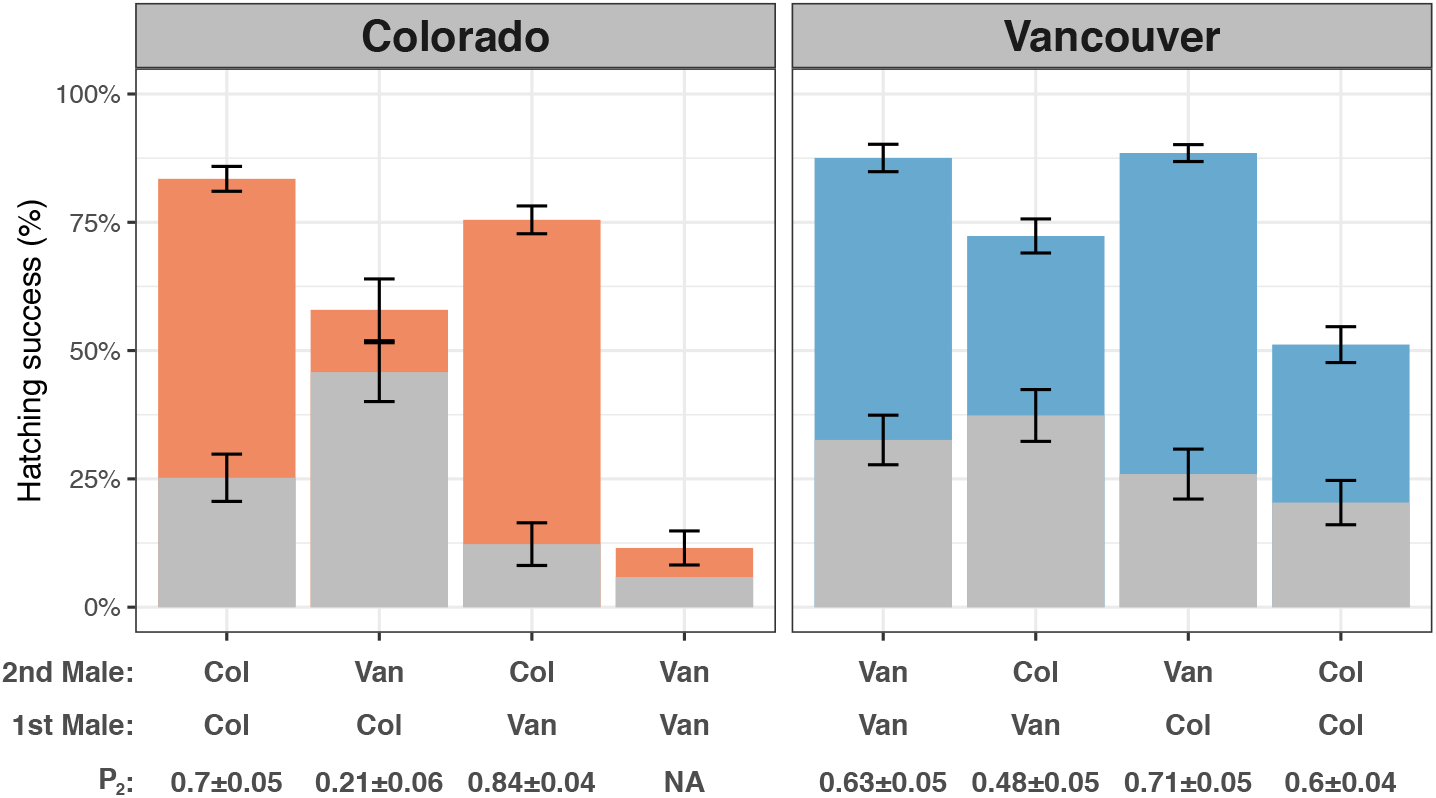
Summary of hatching success and conpopulation sperm precedence (CpSP). Full height of each bar represents mean hatching success (% eggs laid that hatched ± standard error) for females that mated with two nonirradiated males. The upper coloured portion of each bar represents the estimated proportion of offspring sired by the second male to mate (P2 ± standard error), inferred from the irradiated crosses. See Figs. S5 & S6 for results separately. Note: the cross between Colorado females and two Vancouver males was excluded from P2 analysis (see methods and supplementary material). Abbreviations: Col = Colorado, Van = Vancouver.

##### Conpopulation sperm precedence

Females from both populations showed last-male sperm precedence when mating with two within-population males (Colorado [CCC], P2 = 0.70 ± 0.05, n = 28; Vancouver [VVV], P2 = 0.63 ± 0.05, n = 35). However, while there was a significant effect of cross-type on P2 in both Colorado (quasibinomial GLM: F_2,83_ = 65.70, p < 0.001) and Vancouver (quasibinomial GLM: F_3,117_ = 4.93, p = 0.003), only Colorado females showed evidence for CpSP (Fig. 3; Fig. S6). In Colorado female reproductive tracts Colorado males sired the majority of offspring in both the first (CCV, P2 = 0.21 ± 0.06, n = 25) and second (CVC, P2 = 0.84 ± 0.04, n = 34) mating position. In contrast, Vancouver females mating with a Colorado male followed by a Vancouver male showed P2 values that were not significantly different from within-population Vancouver matings (VCV, P2 = 0.71 ± 0.05, n = 27). Vancouver females mating with a Colorado male in the second position used sperm equally from the first and second male (VVC, P2 = 0.48 ± 0.05, n = 31; VCC, P2 = 0.60 ± 0.04, n = 28).

#### Interaction between coevolved and foreign male ejaculates in the female reproductive tract

We found no evidence of an effect of overlapping foreign and coevolved male ejaculates on fertility. Hatching success calculated for the additive effect of two single matings, given estimated P2 values, fell within the range of observed hatching success after a double mating (Fig. S7; Table S2).

### Proxies measuring the intensity of sperm competition within populations

#### Male relative reproductive investment

After dropping the population x soma mass interaction (ANCOVA: F_1,112_ = 0.02, p = 0.884), the reduced model for male relative reproductive investment showed a significant effect of log soma mass on log reproductive tract mass (F_1,113_ = 36.28, p < 0.01) but not of population (F_1,113_ = 0.62, p = 0.433). Log reproductive tract mass increased with log soma mass (Fig. 4a). Thus, we found no difference between populations in male relative investment in reproductive tract tissue.

**Figure 4.**
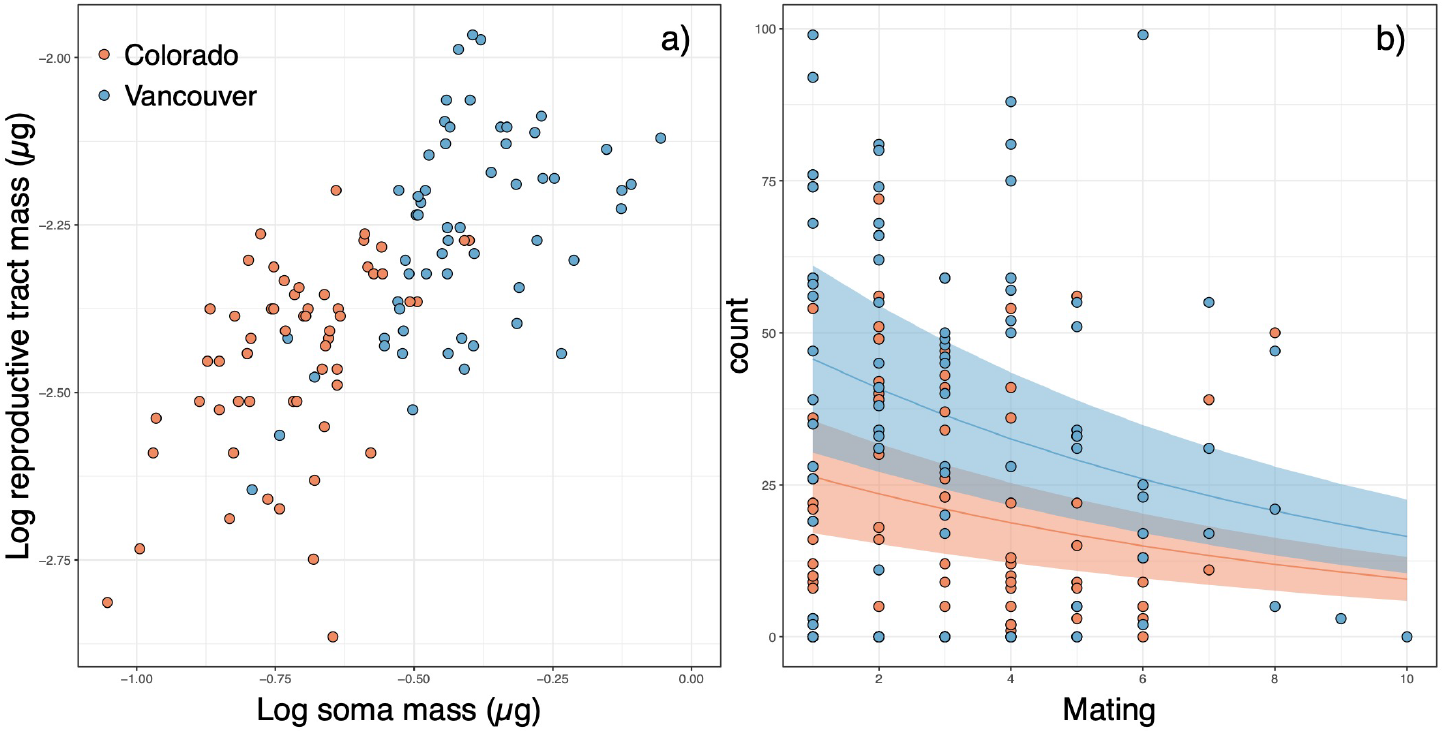
Proxy measures of the intensity of sperm competition experienced by males within populations. Populations did not differ in relative investment in reproductive tract tissue a) or sequential mating capacity and per mating investment b). Colorado, red; Vancouver, blue. Common slopes (±95% confidence intervals) are shown in b). Four outliers were removed in a) and 2 males were excluded in b).

#### Male mating capacity

Populations did not differ in the number of sequential matings that males initiated during the 4-hour observation period (Poisson GLM: χ^2^= 0.01, df = 1, p = 0.939; Colorado = 4.42 ± 0.40, n = 19; Vancouver = 4.47 ± 0.56, n = 19). For male per mating progeny production, there was a significant effect of population (zero-inflated Poisson GLM: χ^2^ = 9.81, df = 1, p = 0.002) and mating number (χ^2^= 146.2, df = 1, p < 0.001). The number of offspring sired declined with mating number (Fig. 4b). The rate of decline was not significantly different between populations (population x mating number interaction, χ^2^= 2.45, df = 1, p = 0.118), suggesting that males from Colorado and Vancouver invested similarly per mating. Vancouver males sired more offspring than Colorado males which resulted in matings within Vancouver producing more offspring than within Colorado overall (quasipoisson GLM: F^1,36^ = 6.40, p = 0.016; Colorado = 93 ± 16, n = 19; Vancouver = 166 ± 25, n = 19). Vancouver females weighed more than Colorado females (t-test, t = −5.81, df = 89.49, p < 0.001; Colorado = 0.77 ± 0.01 μg, n = 60, Vancouver = 0.92 ± 0.02 μg, n = 60). If female body size is indicative of fecundity [58], then this may explain the greater total number of offspring produced in Vancouver crosses.

## DISCUSSION

We used divergent populations of *D. montana* to study the early evolution of conspecific sperm precedence. We performed demographic modelling which revealed a history of divergence with gene flow between North American and Finnish populations. Furthermore, divergence between Colorado and Vancouver began shortly after the split from the ancestral population. We found conpopulation sperm precedence (CpSP) could act as a barrier to gene flow in Colorado females but not Vancouver, showing the same direction of asymmetry to non-competitive PMPZ isolation that we have previously described [35,36]. If the strength of selection acting within populations determines the asymmetry of reproductive isolation, then stronger PMPZ isolation in Colorado is expected to be accompanied by heightened postcopulatory sexual selection. However, we found proxies measuring the intensity of sperm competition faced by males did not differ between populations. Finally, while female multiple mating altered reproductive outcomes, we found no evidence of an interaction between foreign and coevolved ejaculates within the female reproductive tract affecting fertilisation success.

The demographic modelling suggests divergence between Colorado and Vancouver began shortly after the ancestral North American population split from Oulanka. The phylogeographic history of *D. montana* is uncertain, but it is generally assumed to have arisen in eastern Europe or Asia and invaded North America relatively recently [37,59] followed by divergence amongst North American populations during recent ice ages [37]. Our models suggest that the Finnish-North American split occurred 1,750,000 years ago and was followed shortly after by population division within North America (see supplementary material for methods). This is considerably older than previous estimates of timeframe of the North American-Europe division in this species, based on MtDNA sequencing [36]. However, we advise caution when interpreting these estimates due to difficulties in determining how read count data derived from a DNA pool might affect demographic inference and divergence times (see supplementary material). Regardless of actual divergence times, the migration rate estimates suggest gene flow has played a considerable role in the history of divergence between these populations and, importantly, continued following the splits. Colorado and Oulanka also harbored an excess of rare alleles compared to Vancouver, possibly resulting from population expansions after bottlenecks or selective sweeps. Diminished variance in female preferences and male traits has been hypothesised to result in stronger isolation in bottlenecked populations, whereas genetically diverse populations may be more permissive of a greater diversity of genotypes

[60,61]. We might therefore expect Colorado to show lower genetic diversity than Vancouver. However, we found Colorado and Oulanka had larger effective population sizes compared to Vancouver. The most recent genetic analysis identified no fixed SNPs between Colorado and Vancouver [45]. Despite this, the Colorado-Vancouver population pair is the only comparison to show divergence of genes with functional annotations involving reproduction [45] and shows the strongest PMPZ isolation [35,36]. In combination with our results, this shows that specific genes with reproductive function are able to diverge between populations [45] despite the homogenising effects of gene flow reducing genome wide divergence.

We used the irradiated male technique [53] to determine second male paternity share (P2), first showing that last-male sperm precedence occurs for both populations (P2 > 0.63). We subsequently found that CpSP can act as a barrier to gene flow in Colorado because paternity is skewed towards Colorado males when Colorado females mate with Colorado and Vancouver males. In contrast, the Vancouver population did not show evidence of CpSP. When Vancouver females mated with a Colorado male followed by a Vancouver male (VCV) last-male sperm precedence persisted, whereas if the mating order was reversed (VVC) the first and second male sired equal numbers of offspring. Such isolation asymmetries might reflect differences in the strength of postcopulatory sexual selection within populations [18]. However, we found no difference between populations in traits known to evolve in response to the intensity of sperm competition experienced by males [39–41], suggesting sperm competitiveness alone does not predict the strength of PMPZ isolation. It may be that we did not capture the relevant metric of investment. In other species, sperm length and male accessory gland size both respond to manipulation of the strength of sexual selection [9,42,43,62,63]. The pattern of asymmetrical CpSP suggests that Vancouver males can maintain sperm offensiveness against a Colorado male ejaculate in Vancouver female reproductive tracts but cannot maintain a sperm defensive role. In *D. melanogaster* longer and slower sperm are better able to retain representation in the fertilisation set [64]. In *Drosophila* sperm and female sperm storage organ length are highly correlated between populations [62,65] and between species [66] and thought to evolve in response to postcopulatory sexual selection [67]. Sperm-female reproductive tract length coevolution within populations could lead to mismatches between populations resulting in PMPZ isolation.

Finally, we tested whether foreign and coevolved ejaculates interacting in the female reproductive tract altered PMPZ outcomes. Mating with two between-population males produced a similar pattern of PMPZ isolation to that of a single mating, where mating with a foreign male is particularly costly for Colorado females [35,36]. The cost of between-population mating was reduced when females mated with both a within- and between-population male. However, we found no effect of overlapping foreign and coevolved ejaculates on hatching success beyond the expected additive effect of two single matings. This pattern suggests that the ejaculates of the two males act independently, and that cryptic female choice likely plays a more important role than ejaculate x ejaculate interactions in determining PMPZ outcomes in *D. montana*.

To conclude, we found divergence between European and North American *D. montana* populations was followed shortly after by divergence within North America and evolved in the face of gene flow. Conpopulation sperm precedence was stronger in Colorado than Vancouver, showing asymmetry in the same direction as non-competitive PMPZ isolation, suggesting a similar mechanism underlies these phenomena. We show that CSP can evolve between populations, but apparently not in a way predicted by either time since divergence or the strength of sperm competition acting within populations. Male ejaculate x female reproductive tract interactions are complex traits that evolve rapidly and divergently [7,68,69]. The Colorado-Vancouver population pair shows low genome-wide divergence, yet divergence in genes involved in reproduction [45]. The evolutionary processes that can cause such rapid and unpredictable divergence between populations such as sexual coevolution within populations [70] suggests CSP could be present between many taxa where it is currently undocumented. Perhaps such post-mating incompatibilities are a potential example of “mutation-order divergence” [71] and hence would occur sporadically and unpredictably, probably in isolated populations.

## ACKNOWLEDGMENTS

We thank Anneli Hoikkala for providing fly stocks used throughout the experiments, Markus P. Ariaans and Sheila E. Francis for use of the irradiation facility at The University of Sheffield, and Tobit Dehnen and Will Leaning for help with data collection. Sean Stankowski, Roger Butlin, Erin McCullough, and members of the Fitzpatrick lab at Stockholm University provided helpful discussions about the project. Noora Poikela & Maaria Kankare shared access to unpublished data.

---

**Figure S1:**
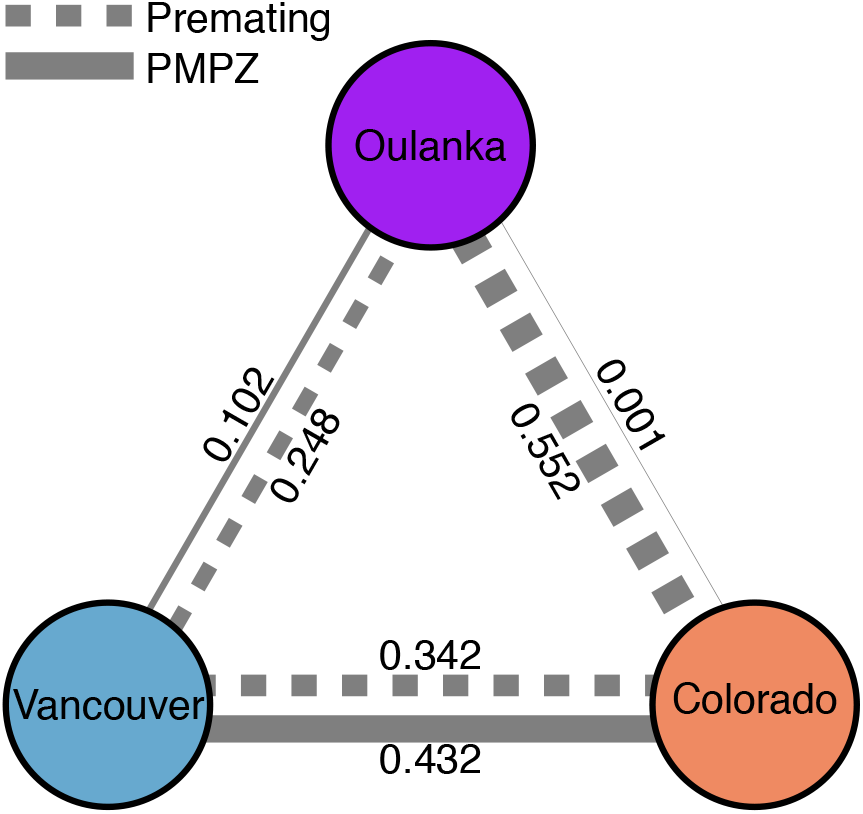
Summary of prezygotic isolation between populations of *Drosophila montana* from Crested Butte, Colorado, USA, Oulanka, Finland, and Vancouver, Canada. Edge weights reflect absolute contributions of premating sexual isolation (dashed) and non-competitive PMPZ isolation (solid) to total reproductive isolation between each pair of populations (from Table 3 in Jennings et al. (2014)).

## Demographic modelling

We tested seven demographic models for the history of divergence between *D. montana* populations from Crested Butte, Colorado, USA, Oulanka, Finland, and Vancouver, Canada (Fig. S2). (1) a no-migration model where migration was not included; (2) a full-symmetrical migration model where migration was symmetrical and present between all populations; (3) an adjacent-migration model where migration was possible between Oulanka and Vancouver or between Colorado and Vancouver, but not between Colorado and Oulanka. Additionally, we included models incorporating periods of isolation falling under two categories: ancient migration models and secondary contact models. Ancient migration models included (4) a model assuming migration ceased at the time of the Colorado-Vancouver split, and (5) a model assuming migration persisted after the Finnish-North American split and the Colorado-Vancouver split but ceased shortly after. For secondary contact models, we included (6) a model with isolation between populations with a recent secondary contact event between Colorado and Vancouver after divergence, and (7) a model assuming migration during the Colorado-Vancouver divergence, with a period of secondary contact beginning after the appearance of both Colorado and Vancouver populations. Migration is assumed to be symmetric for all models.

We fitted all seven models to the 3D-AFS and performed 3 rounds of model optimizations using the Nelder-Mead method (Nelder and Mead 1965). In the first round, minimum (0.01) and maximum (10) parameters were 3-fold perturbed with 10 replicates and a maximum of 5 iterations. The best fitting parameters from the first round were used as starting parameters for the second round, in which parameters were 2-fold perturbed with 10 replicates and a maximum of 5 iterations. Finally, the best fit replicate from the second round was used to produce starting parameters for the third round, in which we ran 15 replicates with parameters perturbed 1-fold with a maximum of 5 iterations.

We estimated divergence times by first solving the equation for *θ*, the population mutation rate, to derive *N_re_f*, the reference effective population size:

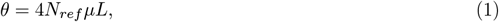

where *μ* is the mutation rate for *D. melanogaster* (2.8×10^-9^) (Keightley et al. 2014) and L, the effective length was calculated by multiplying the total number of SNPs aligned to the reference genome and converted to sync format (31,800,188), by the number of SNPs that entered the analysis divided by the number of SNPs remaining before down-sampling (8,000/860,228). We then converted parameter estimates to real biological units of time, *τi*, using the equation:

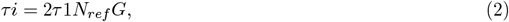

where G, the generation time, was set as one per year based on observations from Aspi et al. (1993) and Throckmorton (1982).

Using equations (1) and (2) we estimated divergence from the ancestral population to have occurred 1,743,164 years ago (T1 + T2), and between Colorado and Vancouver shortly after (T2 = 1,710,355 years ago). We advise caution when interpreting these estimates however. Despite studies showing accurate allele-frequency estimation using Pool-seq (Futschik & Schlötterer 2010; Gautier et al. 2013), it is difficult to appreciate how the use of read count data derived from a DNA pool might affect demographic inference and divergence time estimates. Here, for example, we observe some evidence of an excess of rare variants in our data, though it is unclear whether this represents demographic and/or selective processes, or inaccurate allele frequency estimation caused by inherent statistical challenges associated with Pool-seq. To account for this, we stringently filtered and down-sampled our dataset to retain only reliable sites. Nonetheless, an excess of rare variants may have biased the demographic inference performed here. Additionally, the generation time (G), effect seqeunce length (L) and mutation rate (*μ*) used, whilst these are reasonable assumptions, may have also contributed to over- or underestimation of divergence times.

**Figure S2:**
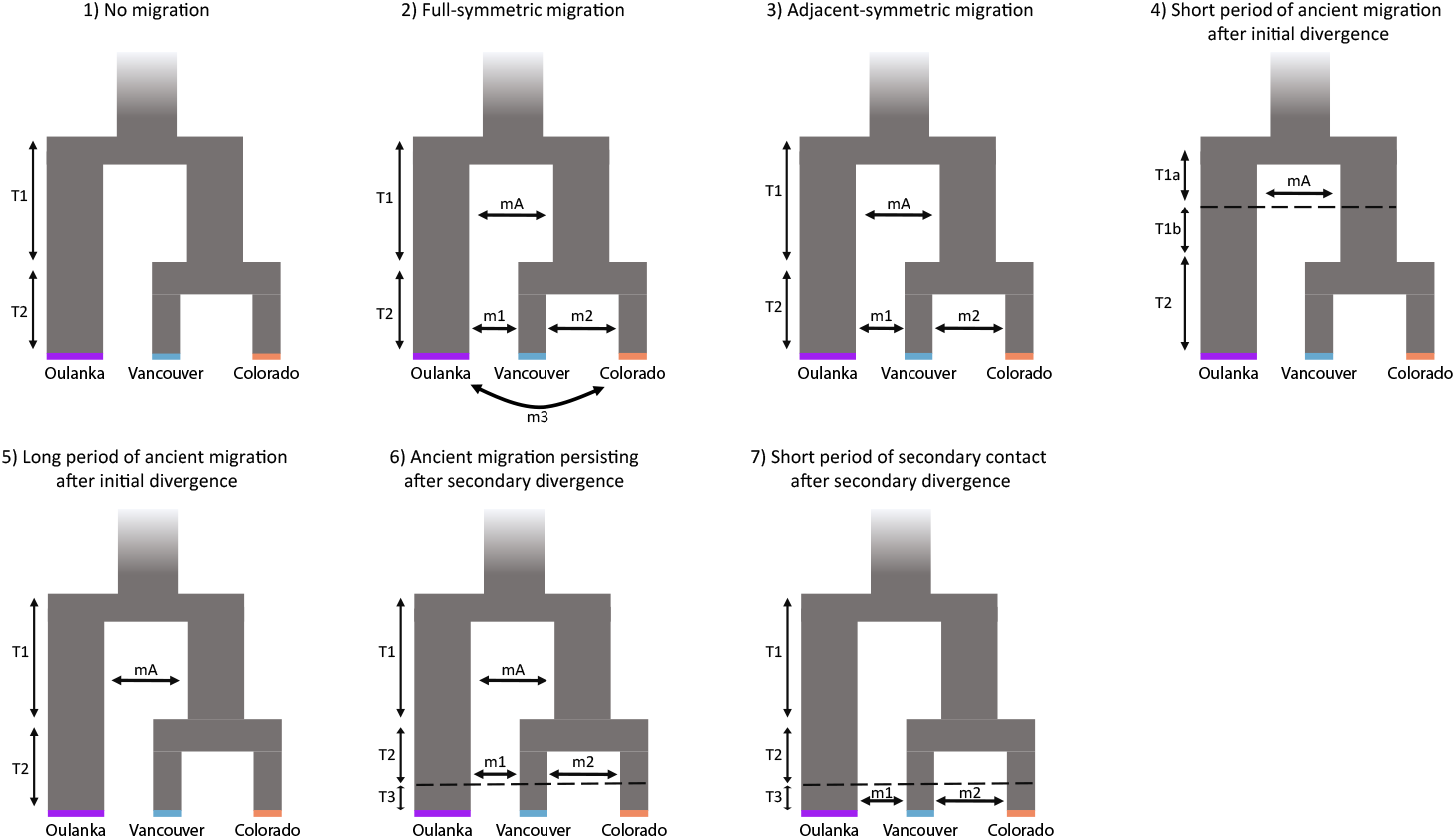
Graphical representation of demographic models tested. mA, ancient migration between Oulanka and precursor North American population; m1, ancient migration between Vancouver and Oulanka; m2, ancient migration between Vancouver and Colorado; m3, ancient migration between Oulanka and Colorado; Initial and secondary divergence are represented by T1(a) and T2, where T1(b) and T3 correspond to the start or end of periods of isolation. Population splits are denoted by T1 (between Oulanka and North America) and T2 (between Vancouver and Colorado). T1(a) and T1(b) denote the start or end of periods of migration following the split between Oulanka and North American populations. T3 denotes start or end of periods of migration following the split between Vancouver and Colorado.

**Figure S3:**
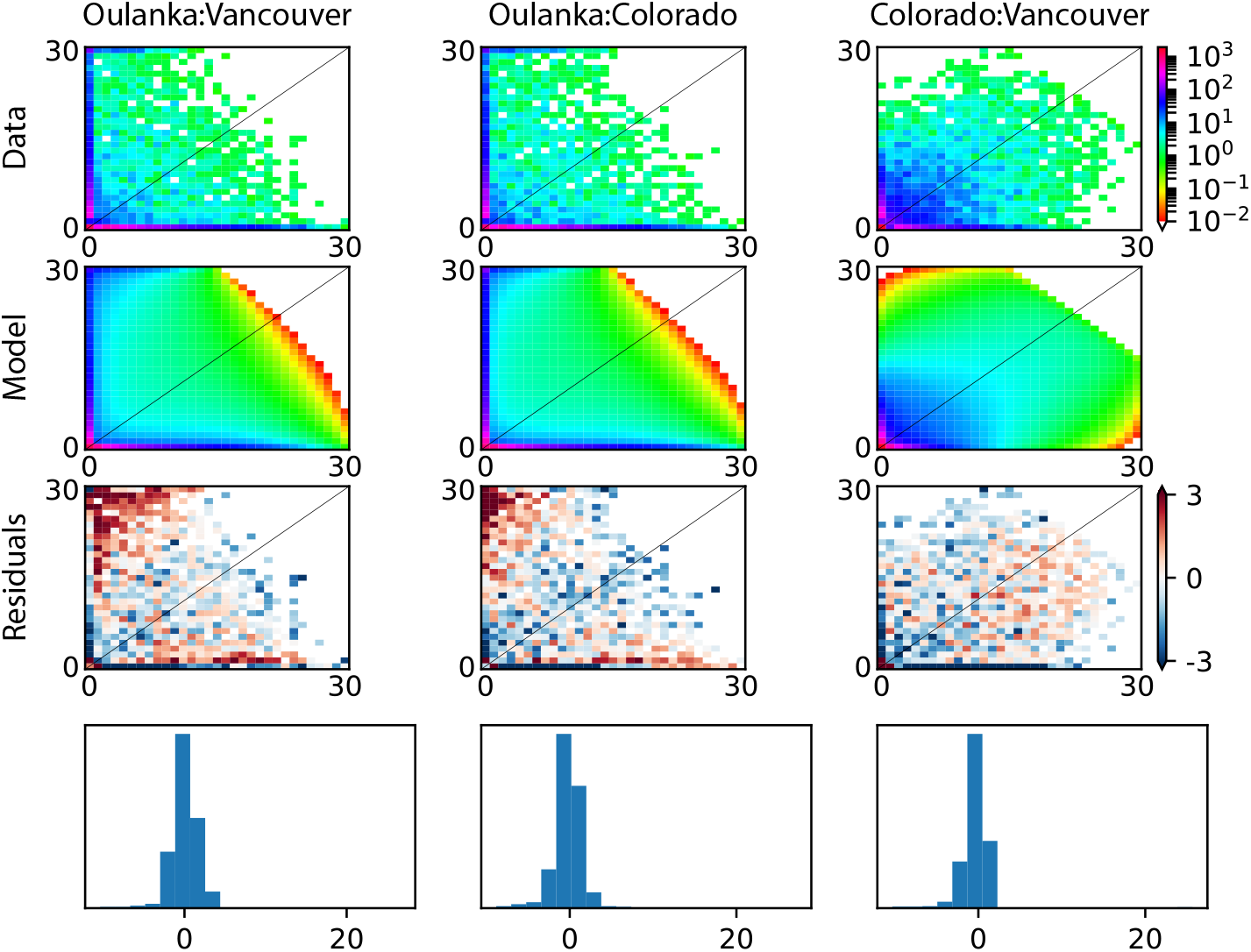
Three-dimensional allele-frequency spectra (3D-AFS) for the best fitting model; (3) Adjacent-migration model. Top: data; middle top: model fits; middle bottom: residuals; bottom: histogram of residuals.

## Measures of postmating prezygotic reproductive isolation

### Postmating prezygotic isolation of virgin females: non-competitive gametic isolation (NCGI)

**Figure S4:**
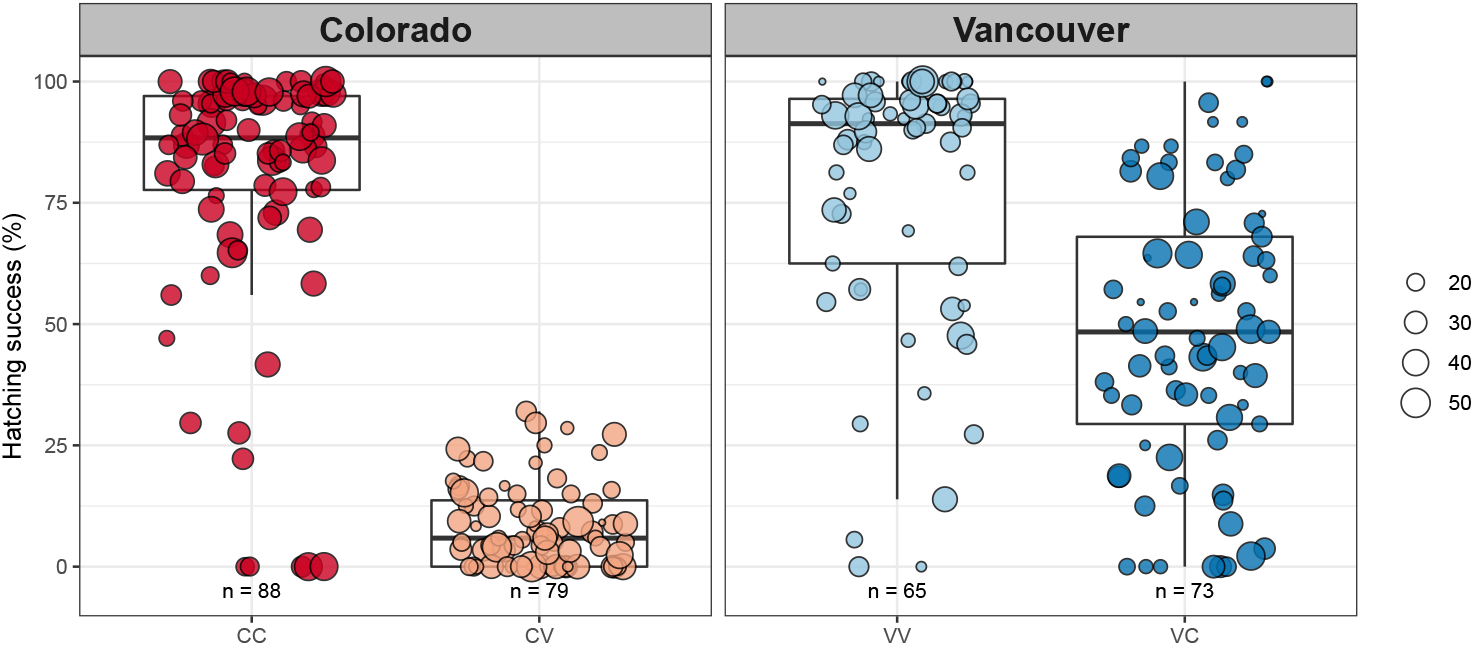
Hatching success (% eggs laid that hatched) after the first mating for females that mated with nonirradiated males. Points are observations and size show the number of eggs laid (i.e. weights). Cross-type denotes female population followed by first and second male. C, Colorado; V, Vancouver.

### Postmating prezygotic isolation of non-virgin females

**Figure S5:**
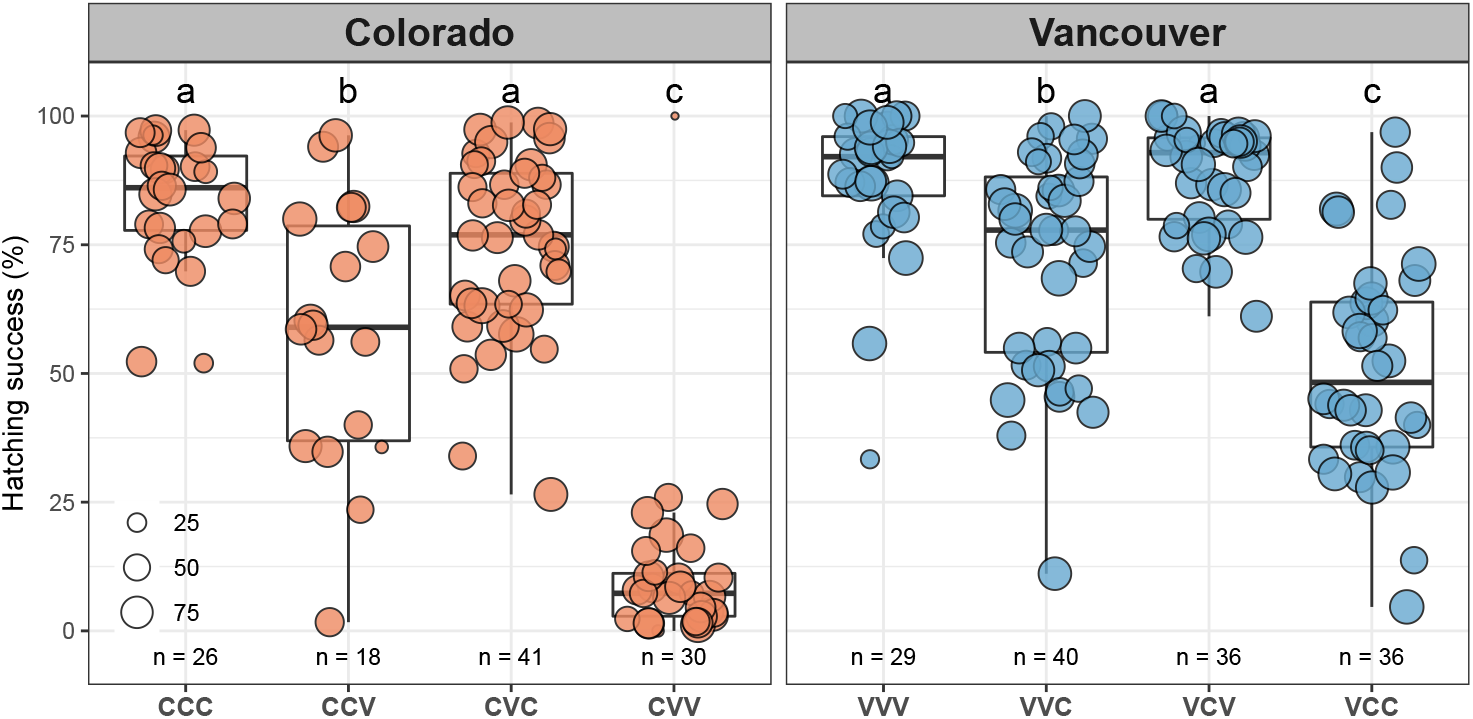
Hatching success (% eggs laid that hatched) after the second mating for females that mated with nonirradiated males. Points are observations and size show the number of eggs laid (i.e. weights). Cross-type denotes female population followed by first and second male. C, Colorado; V, Vancouver.

### Con-population sperm precedence (CpSP)

We assessed male offensive paternity share (*P*_2_) using the irradiated male technique (Boorman and Parker 1976). Virgin males were irradiated with a 100Gy dose of gamma radiation (dose rate 189.2 rads min^-1^, ^137^Cs source), rendering males 100% sterile, less than 24 hours before mating and housed individually overnight in food vials. To assess the level of fertility after a double mating, ‘control’ treatments consisted of mating females with two nonirradiated males in all possible crossing combinations with males from Colorado and/or Vancouver. We also included additional data for these control crosses collected in a pilot study (n = 30 per cross-type). To assess the efficacy of the irradiation technique females were mated with two irradiated males, both from Colorado or Vancouver. Experimental treatments consisted of all possible crossing combinations between females and males from Colorado and Vancouver, with either the first or second male irradiated. The proportion of eggs fertilised by the irradiated male after the second mating, *P_R_*, was calculated using equation (1) from Boorman and Parker (1976):

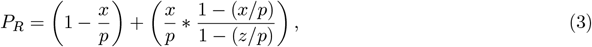

where *x* is the observed proportion of developing eggs after a double mating from the second oviposition plate, *p* is the level of fertility observed in a double mating for a given ‘control’ cross-type, and *z* is the level of fertility observed in the double irradiated cross-type. For our calculation of *P_R_* we chose the maximum value of *p* observed (rather than the mean) to capture the full potential fertility of between-population crosses. As *z* = 0 in this case, the equation can be simplified to *P_R_* = 1 — *x/p*. Therefore, if the irradiated male mates first, then the proportion of eggs fertilised by the second male, *P*_2_ = *x/p*. If the irradiated male mates second then *P*_2_ = *P_R_* (Boorman and Parker 1976). The total number of eggs laid by each female after the second mating was multiplied by the calculated *P*2 value and rounded to a whole number to give the estimated number of offspring sired by the second male. The remaining number of eggs expected to hatch were assigned to the first male.

As few eggs laid hatch when Colorado females mate with Vancouver males (CV hatching success = 0.084 ± 0.01 [mean ± standard error], n = 79, Fig. S4; CVV hatching success = 0.115 ± 0.033, n = 30, Fig. S5), we could not estimate differences in *P*_2_ for the CVV cross-type which was subsequently excluded from *P*_2_ analyses (Fig. S6).

**Table 1:** Mean *P*2 with standard errors. Value calculated for CVV cross is included for completeness. C, Colorado; V, Vancouver. N = sample size.

| Cross-type | N | $P_2$ | Std. Err. |
| --- | --- | --- | --- |
| CCC | 28 | 0.70 | 0.05 |
| CCV | 25 | 0.21 | 0.06 |
| CVC | 33 | 0.84 | 0.04 |
| CVV | 25 | 0.49 | 0.09 |
| VVV | 35 | 0.63 | 0.05 |
| VVC | 31 | 0.48 | 0.05 |
| VCV | 27 | 0.71 | 0.05 |
| VCC | 28 | 0.60 | 0.04 |

### Irradiation order effects on *P*_2_

In Colorado there was a significant effect of irradiation treatment (*F_1,82_* = 20.41, p < 0.001), and the cross-type x irradiation interaction on *P*_2_ (*F*_2,80_ = 14.12, p < 0.001). Irradiated males mating in the second position had a greater *P*2 than irradiated males mating in the first position in the CCV and CVC cross-types, but not the CCC cross-type (Fig. S6). In Vancouver the irradiation main effect was not significant (*F*_1,116_ = 1.24, p = 0.267) but there was a significant cross-type x irradiation interaction (*F_3,113_* = 7.37, p < 0.001). In the VVV cross-type irradiated males in the second position had lower *P_2_*, whereas in the other Vancouver female cross-types irradiated males in the second position had greater *P_2_* (Fig. S6).

**Figure S6:**
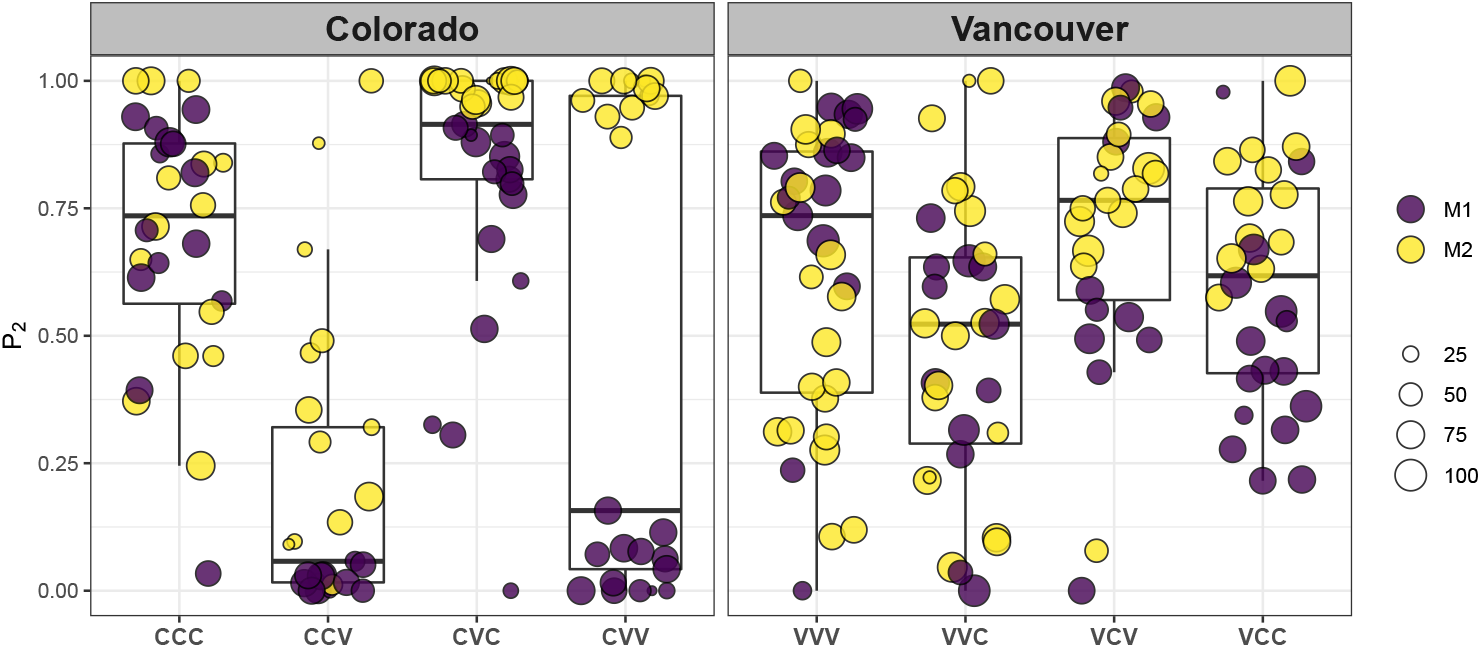
Proportion of offspring sired by the second male to mate (*P*_2_), where either the first (M1, purple) or second (M2, yellow) male to mate was irradiated. Cross-type denotes female population followed by first and second male mate. C, Colorado; V, Vancouver. Points are observations and size show the number of eggs laid (i.e. weights).

### Interaction between coevolved and foreign male ejaculates in the female reproductive tract

We tested whether observed hatching success rates after a double mating in the nonirradiated ‘control’ crosses differed from the expected additive effect of two single matings, *H_total_*, using the following equation:

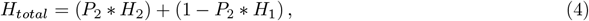

where *P*_2_ is the mean proportion of offspring sired by the second male in a given cross-type and *H*ļ and *H*_2_ are the mean hatching success rates after a single mating for a female mated with a male from the first, and second, population denoted in a cross-type, respectively (Table 2).

**Table 2:** Values used to calculate *H_total_*, the expected additive effect of two single matings using equation (4). Cross-type denotes female population followed by first and second male mate. C, Colorado; V, Vancouver.

| Cross-type | $P_2$ | $H_{obs}$ | $H_1$ | $H_2$ | RHS | LHS | $H_{total}$ | $\Delta$ |
| --- | --- | --- | --- | --- | --- | --- | --- | --- |
| CCC | 0.70 | 0.83 | 0.80 | 0.80 | 0.24 | 0.56 | 0.80 | 0.03 |
| CCV | 0.21 | 0.58 | 0.80 | 0.08 | 0.63 | 0.02 | 0.65 | -0.07 |
| CVC | 0.84 | 0.75 | 0.08 | 0.80 | 0.01 | 0.67 | 0.68 | 0.07 |
| CVV | 0.49 | 0.12 | 0.08 | 0.08 | 0.04 | 0.04 | 0.08 | 0.03 |
| VVV | 0.63 | 0.88 | 0.78 | 0.78 | 0.29 | 0.49 | 0.78 | 0.09 |
| VVC | 0.48 | 0.72 | 0.78 | 0.48 | 0.41 | 0.23 | 0.64 | 0.09 |
| VCV | 0.71 | 0.88 | 0.48 | 0.78 | 0.14 | 0.55 | 0.69 | 0.19 |
| VCC | 0.60 | 0.51 | 0.48 | 0.48 | 0.19 | 0.29 | 0.48 | 0.03 |

$P_2$ , mean second male paternity share; $H_{obs}$ , mean observed hatching success for females that mated with two nonirradiated males; $H_1$ , mean hatching success from a single mating expected for females mating with first male; $H_2$ , mean hatching success from a single mating expected for females mating with second male; RHS, $1 - P_2 * H_1$ ; LHS, $P_2 * H_2$ ; $H_{total}$ , LHS + RHS; $\Delta = H_{obs} - H_{total}$

**Figure S7:**
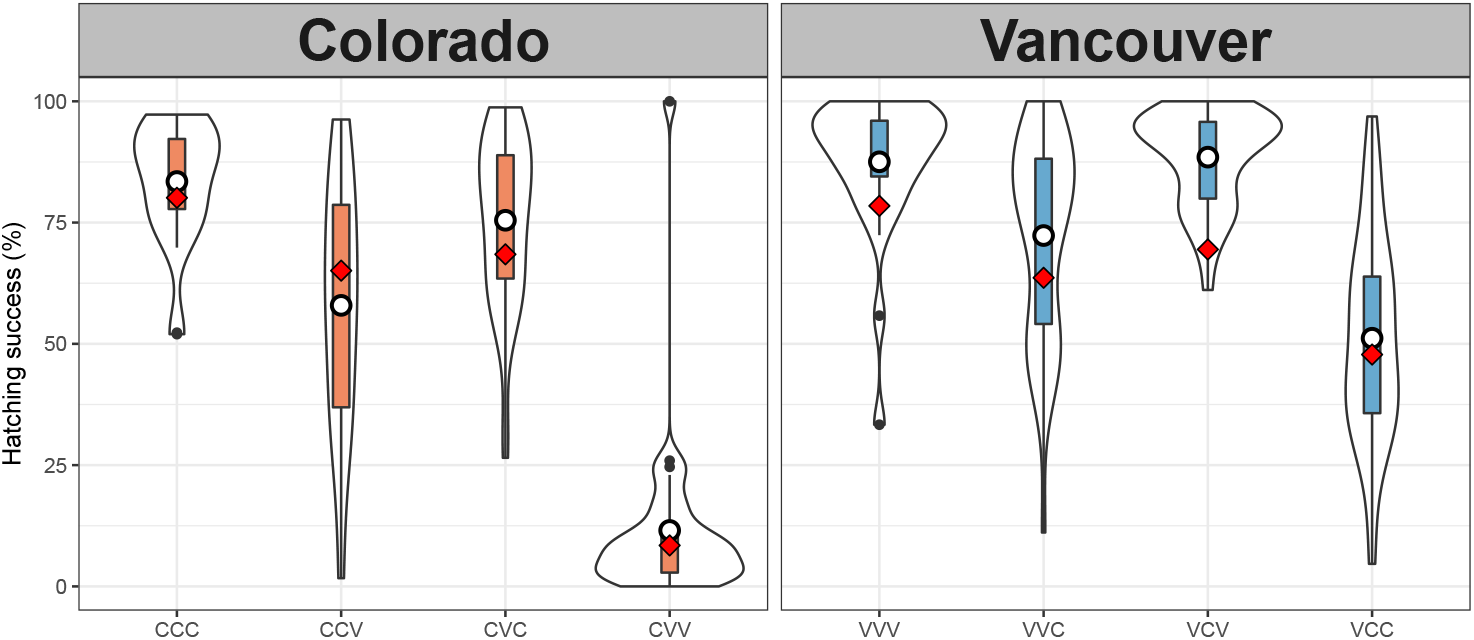
Effects of overlapping foreign and coevolved ejaculates on fertilisation success. White points show *H_obs_*, mean hatching succcess (% eggs laid that hatched) for females that mated with two nonirradiated males (data same as Fig. S5), red diamonds show *H_total_*, the expected additive hatching success of two single matings for a female mated with a male from the first and second population denoted in a cross-type, calculated given the estimated paternity share of each male (Table 2). Cross-type denotes female population followed by first and second male mate. C, Colorado; V, Vancouver.

## Proxies for the intensity of sperm competition faced by males

### Male relative reproductive investment

We tested for differences between populations in male reproductive mass investment using analysis of covariance (ANCOVA) (Tomkins and Simmons 2002). Log transformed dry reproductive tract mass was regressed against log transformed dry soma mass (body mass – reproductive tract mass) and the soma mass x population interaction. Colorado males weighed significantly less than Vancouver males on average (Welch Two Sample t-test, t = −9.84, df = 96.45, p < 0.001; Colorado = 0.58 ± 0.01 mg [mean ± standard error], n = 60, Vancouver = 0.77 ± 0.02 mg, n = 60; Fig. S8a). Colorado males reproductive tracts weighed significantly less than Vancouver males on average (Welch Two Sample t-test, t = −6.47, df = 114.14, p < 0.001; Colorado = 0.09 ± 0.002 mg, n = 60, Vancouver = 0.11 ± 0.002 mg, n = 60; Fig. S8c). The results of the ANCOVA revealed the population x log soma mass interaction was not significant (ANCOVA: *F_1,116_* = 0.995, p = 0.321) and so was dropped from the model. The reduced model showed a significant effect of population (ANCOVA: *F*_1,117_ = 7.74, p = 0.006) and log soma mass (ANCOVA: *F*_1,117_ = 8.63, p = 0.004) on reproductive mass. Log reproductive tract mass increased with log soma mass and Vancouver males had a higher intercept (Fig. S8b). However, visual inspection of model diagnostic plots revealed four consistent outliers (Fig. S9). One Vancouver male had an unusually small body mass which resulted in high leverage (Fig. S8a). Three males (2 Colorado, 1 Vancouver) had relatively small reproductive tract masses (Fig. S8c).

**Figure S8:**
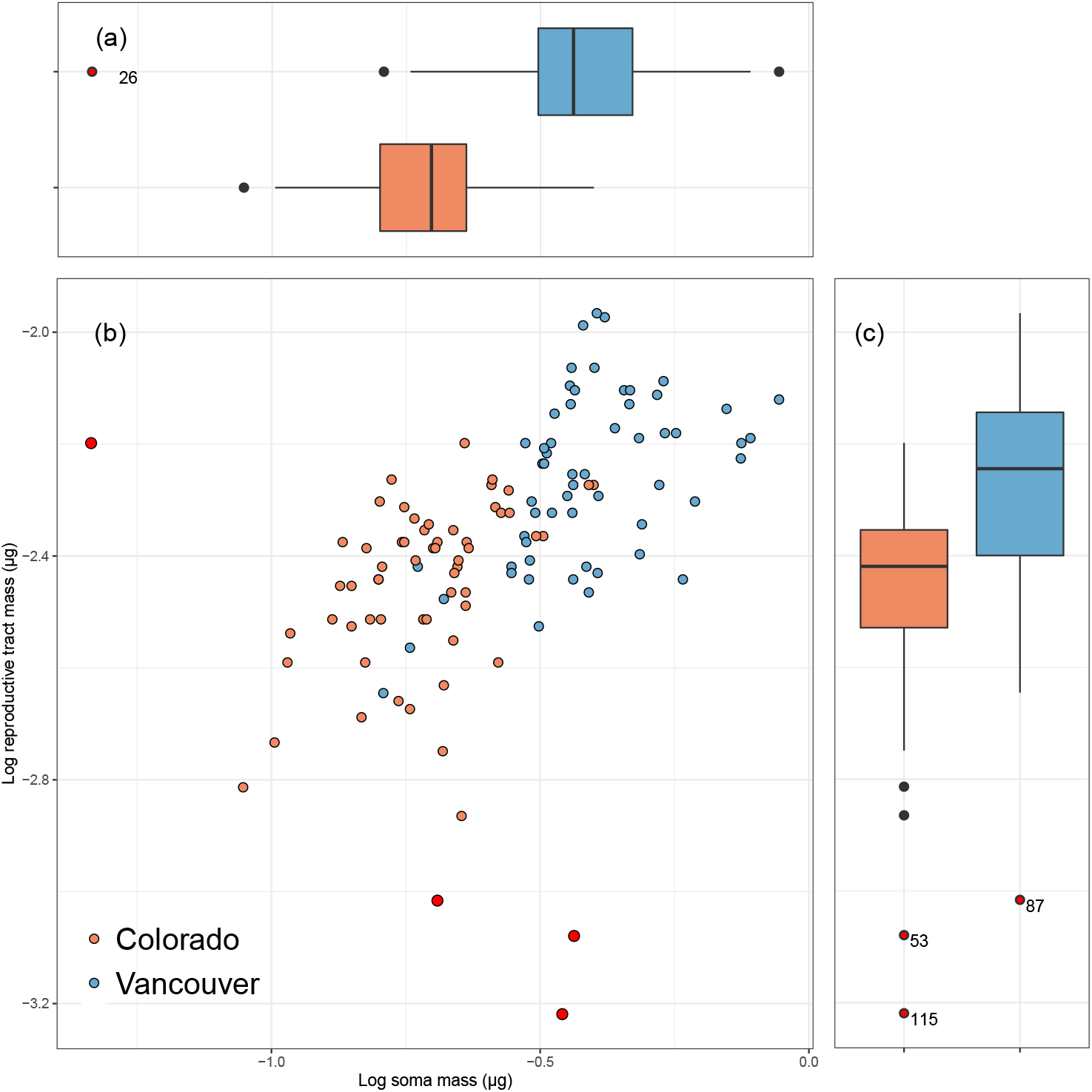
Log reproductive tract mass increased with log soma mass (ANCOVA: common slope, *β* = 0.520 ± 0.086, t = 6.023, p < 0.001). Red, Colorado; Blue, Vancouver. Outliers removed for analysis (2 Colorado, 2 Vancouver) are numbered and shown in red.

**Figure S9:**
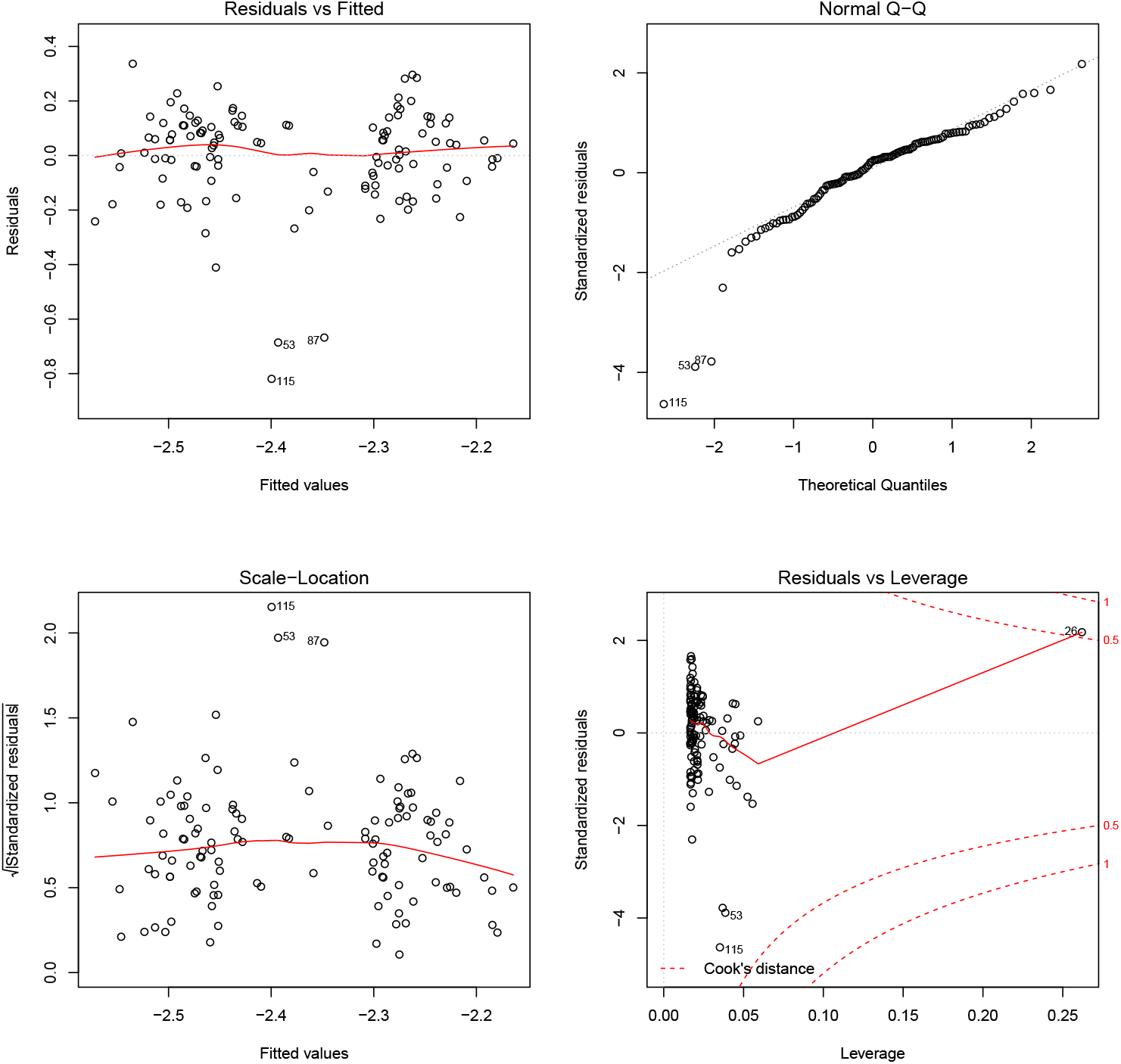
Model diagnostic plots for ANCOVA after dropping the population x soma mass interaction including all data. Outliers are numbered as in Fig. S8.

**Figure S10:**
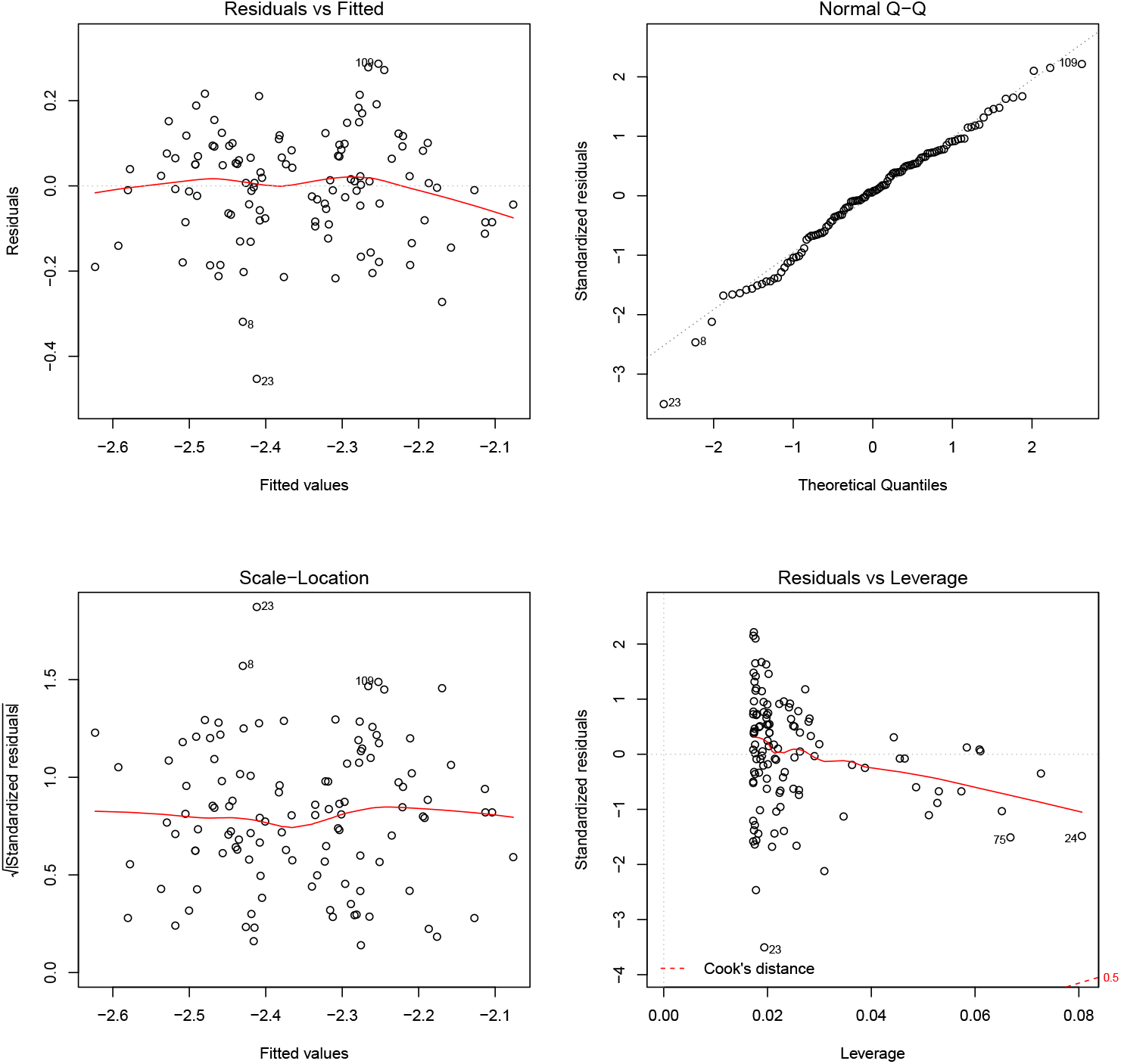
Model diagnostic plots for ANCOVA after dropping the population x soma mass interaction model after excluding outliers.

After removing the four outliers, Colorado males still weighed significantly less than Vancouver males on average (Welch Two Sample t-test, t = −11.36, df = 98.8, p < 0.001) and Colorado reproductive tracts weighed significantly less than Vancouver males on average (Welch Two Sample t-test, t = −6.91, df = 106.27, p < 0.001). Results of the updated ANCOVA model again revealed that the population x log soma mass interaction was not significant (ANCOVA: *F*_1,112_ = 0.02, p = 0.884), and was subsequently dropped from the model. Visual inspection of model diagnostic plots showed no significant outliers (Fig. S10).

